# PepNN: a deep attention model for the identification of peptide binding sites

**DOI:** 10.1101/2021.01.10.426132

**Authors:** Osama Abdin, Satra Nim, Han Wen, Philip M. Kim

## Abstract

Protein-peptide interactions play a fundamental role in facilitating many cellular processes, but remain underexplored experimentally and difficult to model computationally. Here, we present PepNN-Struct and PepNN-Seq, structure and sequence-based approaches for the prediction of peptide binding sites on a protein given the sequence of a peptide ligand. A main difficulty for the prediction of peptide-protein interactions is the flexibility of peptides and their tendency to undergo conformational changes upon binding. To account for this behaviour, we developed a novel reciprocal attention module that simultaneously updates the encodings of peptide and protein residues and explicitly enforces the symmetry in the updates, allowing for information flow and reflecting the biochemical reality of conformational changes in the peptide. PepNN additionally makes use of modern graph neural network layers that are effective at learning representations of molecular structure. Finally, to compensate for the scarcity of peptide-protein complex structural information, we make use of available protein-protein complex and protein sequence information through a series of transfer learning steps. PepNN-Struct achieves state-of-the-art performance on the task of identifying peptide binding sites, with a ROC AUC of 0.893 and an MCC of 0.483 on an independent test set. Beyond prediction of binding sites on proteins with a known peptide ligand, we also show that the developed models make reasonable peptide-agnostic predictions, allowing for the identification of novel peptide binding proteins.

## Introduction

Interactions between proteins and peptides are critical for a variety of biological processes. A large fraction of protein-protein interactions are mediated by the binding of intracellular peptide recognition modules (PRMs) to linear segments in other proteins^1^. Moreover, peptide ligands binding to extracellular receptors have important functions^2^. In total, it is estimated that there are roughly 10^4^ human proteins that contain at least one PRM^3^ and that there are over 10^6^ peptide motifs encoded in the human proteome^1^. Disruption of these interactions and their regulation can consequently result in disease; for instance, many proteins with PRMs harbor oncogenic mutations^4^. It has also been shown that viral proteins encode peptidic motifs that can potentially be used to hijack host machinery during infection^5^.

In the absence of ample experimental data including solved structures, gaining molecular insight into these interactions and their associated disease states is contingent on the ability to model peptide binding computationally. This has been a difficult problem that has traditionally been approached with peptide-protein docking^6^. One widely used peptide docking tool is FlexPepDock, a Rosetta protocol that refines coarse-grain peptide-protein conformations by sampling from the degrees of freedom within a peptide^7^. In general, benchmarking studies have shown that peptide docking approaches often fail to accurately identify the native complex conformation^8–10^, indicating that this problem remains unsolved; current approaches are limited by the high flexibility of peptides as well the inherent error of scoring heuristics^6^. Machine learning approaches provide potential alternatives to docking, as they can sidestep the issue of explicit enumeration of conformational space and can learn scoring metrics directly from the data.

A number of machine learning approaches have been applied to the problem of predicting the binding sites of peptides with varying amounts of success^11–16^. More recently, deep learning approaches have resulted in large improvements in many areas, including in the domains of protein and structural biology^17^. However, no such model has been developed for the identification of peptide binding sites.

Here, we sought to develop a novel deep learning architecture to improve upon existing approaches. In particular, we develop an architecture that is partially inspired by the Transformer, a model that primarily consists of repeated multi-head attention modules^18^. These modules are effective at learning long-range dependencies in sequence inputs and have been successfully adapted to graph inputs^19^. Graph neural networks in general have had success on various related problems, including protein design^19,20^. Importantly, we build upon attention to develop reciprocal attention, a variant that updates two input encodings based on dependencies between the two encodings, while maintaining symmetry in the updates. This novel type of architecture updates the embedding of both residues participating in an interaction simultaneously, and reflects the fact that the conformation of a bound peptide depends on the interacting protein target^21^.

One significant hurdle to the development of deep learning approaches for the modelling of peptide-protein complexes has been the paucity of available training data. To overcome this problem, we exploit available protein-protein complex information, thereby adding an order of magnitude more training data. The “hot segment” paradigm of protein-protein interaction suggests that the interaction between two proteins can be mediated by a linear segment in one protein that contributes to the majority of the interface energy^22^. Complexes of protein fragments with receptors thus represent a natural source of data for model pre-training. In addition, the idea of pre-training contextualized language models has recently been adapted to protein biology for the purpose of generating meaningful representations of protein sequences^23,24^. The success of these approaches provides an opportunity to develop a strictly sequence based peptide binding site predictor.

In this study, we integrate the use of contextualized-language models, available protein-protein complex data, and a task-specific attention-based architecture, to develop parallel models for both structure and sequence-based peptide binding site prediction: PepNN-Struct and PepNN-Seq. Comparison to existing approaches reveals that our models perform better in most cases. We also show that the developed models can make reasonable peptide-agnostic predictions, allowing for their use for the identification of novel peptide binding sites.

## Results

### Parallel models for structure and sequence-based peptide binding site prediction

PepNN takes as input a representation of a protein as well as a peptide sequence, and outputs residue-wise scores representing the confidence that a particular residue is part of a peptide binding site (Fig 1a-b). The PepNN-Seq and PepNN-Struct architectures are based in part on the Transformer and a graph variant of the Transformer^18,19^. PepNN-Struct makes use of graph attention layers to learn from context within an input protein (Fig 1a). PepNN-Seq generates predictions based solely on the input protein and peptide sequences (Fig 1b).

**Figure 1:**
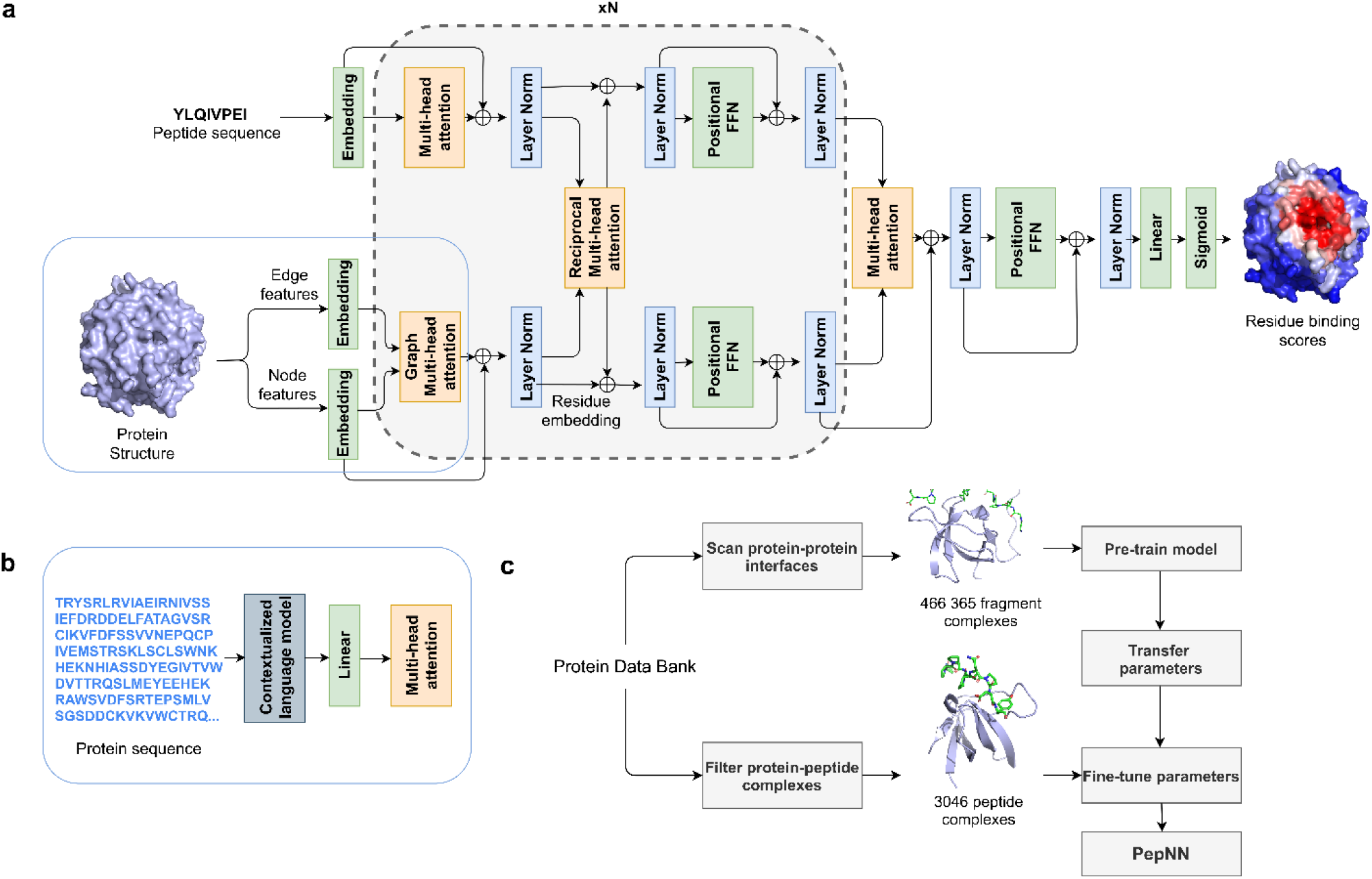
Model architecture and training procedure. **a)** Attention layers are indicated with orange, normalization layers are indicated with blue and simple transformation layers are indicated with green. **b)** Input layers for PepNN-Seq. **c)** Transfer learning pipeline used for model training.

PepNN differs from conventional Transformers in that it does not follow an encoder-decoder architecture. Encoding the peptide sequence independently from the protein representation would implicitly assume that all information about the peptide is contained within its sequence and this assumption is not concordant with the fact that many disordered regions undergo conformational changes upon protein binding^21^. In other words, a peptide’s sequence is insufficient by itself to determine its bound conformation. In fact, the same peptide can adopt different conformations when bound to different partners. To explicitly reflect this process, we introduce multi-head reciprocal attention layers, a novel attention-based module that simultaneously updates the peptide and protein embeddings while ensuring that the unnormalized attention values from protein to peptide residues are equal to the unnormalized attention values in the other direction. This ensures that the protein residues involved in binding have influence on the peptide residues and vice versa, representing the physical reality of the peptide-protein binding process. The exact model hyperparameters were determined using random search (see Methods) and we compared the performance of the model to a graph Transformer with the same hyperparameters on the preliminary task of identifying the binding sites of protein fragments. We found that the reciprocal attention variant outperforms the graph Transformer in its capacity to accurately identify fragment binding sites (Fig S1).

### Transfer learning results in large improvements in model performance

We used transfer learning in two ways to improve model performance. The first was to pretrain the model on a large protein fragment-protein complex dataset before fine-tuning with a smaller dataset of peptide-protein complexes (Fig 1c). To generate the fragment dataset, we scanned all protein-protein complex interfaces in the PDB with the PeptiDerive Rosetta protocol^25^ to identify protein fragments of length 5-25 amino acids that contribute to a large portion of the complex interface energy (Fig S2). These fragment-protein complexes were filtered based on their estimated interface energy as well as the buried surface area to ensure that they had binding properties that were reasonably close to that of peptide-protein complexes. The second application of transfer learning was the use a pre-trained contextualized language model, ProtBert^23^, to embed protein sequences. These high dimensional, information-rich, embeddings were used as input to PepNN-Seq (Fig 1b).

To evaluate the impact of transfer learning on model performance, we trained PepNN-Struct and PepNN-Seq using different procedures. Pre-training PepNN-Struct resulted in significant improvement over models trained on only the fragment or peptide complex dataset, both in terms of over all binding residue prediction, and in terms of prediction for individual proteins (Fig 2a, b). Model predictions on the Bro domain of HD-PTP demonstrate this difference in performance, as only the pre-trained variant of the model correctly predicts the peptide binding site (Fig 2c). This example furthermore illustrates that the pre-training step helps bring the parameters closer to an optimum for general peptide binding site prediction, rather than improving performance solely on examples that match patterns seen in the fragment-complex dataset.

**Figure 2:**
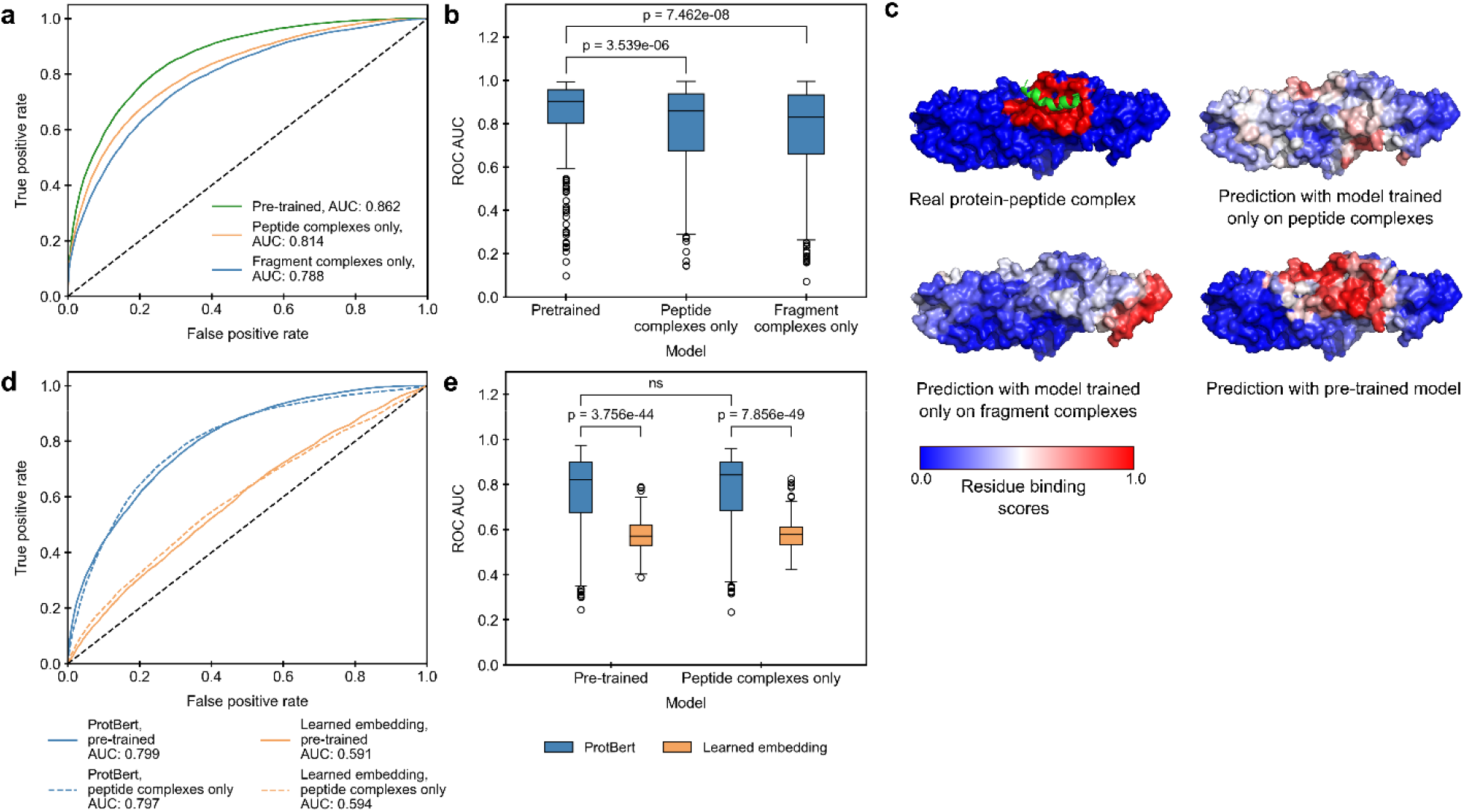
Impact of transfer learning on model performance on the peptide complex validation dataset. **a)** ROC curves on all residues in the dataset using predictions from PepNN-Struct trained on different datasets. **b)** Comparison of the distribution of ROC AUCs on different input proteins using predictions from PepNN-Struct with different training procedures and sequence embeddings (Wilcoxon signed-rank test). **c)** Predictions of the binding site of the Bro domain of HD-PTP (PDB code 5CRV) using PepNN-Struct trained on different datasets. **d)** ROC curves on all residues in the dataset using predictions from the sequence model with different training procedures and sequence embeddings. **e)** Comparison of the distribution of ROC AUCs on different input proteins using predictions from PepNN-Seq trained on different datasets (Wilcoxon signed-rank test).

Embedding protein sequences with ProtBert resulted in large performance improvements over learned embedding parameters for PepNN-Seq (Fig 2d, e). Interestingly, pretraining on the fragment complexes did not have a large impact on PepNN-Seq performance (Fig 2b, d). This suggests that pre-training on the fragment complexes allows PepNN-Struct to learn reasonable protein embeddings while the use of a pre-trained contextualized language model is sufficient for the generation of reasonable embeddings in the case of PepNN-Seq.

### PepNN achieves state-of-the-art performance on peptide binding site prediction

We initially evaluated the developed models on an independent test set derived from the peptide complex dataset. Unsurprisingly, we found that PepNN-Struct outperforms PepNN-Seq (Table 1). We additionally ran the sequence-based PBRpredict-Suite model on this test dataset^16^. All three variants of this model performed worse than PepNN on this dataset (Table 1) and notably, the observed performance was drastically lower than the performance reported in the original publication. This could potentially be due to the fact a smoothing approach was used to annotate binding sites in the PBRpredict-Suite study^16^, while binding site residues annotations were made based only on distance to peptide residues in this study.

**Table 1:** Comparison of the developed model to existing approaches

| Test dataset | Training dataset size | Model | ROC AUC | MCC |
| --- | --- | --- | --- | --- |
| TS305 | 2394 | PepNN-Struct | **0.893** | **0.483** |
|  |  | PepNN-Seq | 0.859 | 0.401 |
|  | 475 | PBRpredict-flexible <sup>16</sup> | 0.653 | 0.139 |
|  |  | PBRpredict-moderate <sup>16</sup> | 0.620 | 0.127 |
|  |  | PBRpredict-strict <sup>16</sup> | 0.598 | 0.100 |
| TS251 | 251 | PepNN-Struct | **0.817** | **0.370** |
|  |  | PepNN-Seq | 0.758 | 0.278 |
|  |  | Interpep <sup>11</sup> | **0.793** | --- |
| TS639 | 640 | PepNN-Struct | **0.838** | 0.301 |
|  |  | PepNN-Seq | 0.792 | 0.251 |
|  |  | PepBind <sup>12</sup> | 0.767 | **0.348** |
| TS125 | 640 | PepNN-Struct | **0.841** | 0.321 |
|  |  | PepNN-Seq | 0.805 | 0.278 |
|  |  | PepBind <sup>12</sup> | 0.793 | **0.372** |
|  | 1156 | SPRINT-Str <sup>41</sup> | 0.780 | 0.290 |
|  | 1199 | SPRINT-Seq <sup>13</sup> | 0.680 | 0.200 |
|  | 1004 | Visual <sup>15</sup> | 0.730 | 0.170 |

Most other existing approaches lack programmatic access and a portion rely on alignments to reference datasets that overlap with the test set. We hence used values reported in the literature for comparison. To ensure an unbiased comparison, the model was re-trained on the training datasets used in different studies prior to comparison on their test sets. In all cases, PepNN-Struct largely outperforms existing approaches in terms of ROC AUC (Table 1). In most cases, PepNN-Seq also outperforms existing approaches by this metric. PepNN does, however, perform worse in terms of MCC in a couple of cases, suggesting that there exist thresholds at which the models do not perform was well as the PepBind approach, despite having more robust performance at different prediction thresholds. It is worth noting that the training datasets used in other studies were substantially smaller and thus training on them resulted in lower performance of our models overall (Table 1). This was because the datasets used in other studies are relatively outdated and a smaller portion of the available data was used for training.

### Peptide-agnostic prediction allows the identification of putative novel peptide binding proteins

To quantify the extent to which the model relies on information from the protein when making predictions, we tested the ability of PepNN-Struct and PepNN-Seq to predict peptide binding sites using random length poly-glycine peptides as input sequences. While the models did perform better when given the native peptide sequence than with a poly-glycine sequence (p-value < 2.2e-16 for both PepNN-Struct and PepNN-Seq, DeLong test), there was only a small overall decrease in the ROC AUC when a poly-glycine peptide was given (Fig 3a, b). Comparing the probabilities that the model assigns to different residues shows that in both the case of PepNN-Struct and PepNN-Seq, providing the native peptide increases the model’s confidence when predicting binding residues (Fig S3). Providing the native peptide sequence is thus important for reducing false negatives. Overall, these results suggest that while providing a known peptide can increase model accuracy, the model can make reasonable peptide-agnostic predictions and could potentially be used to identify novel peptide binders.

**Figure 3:**
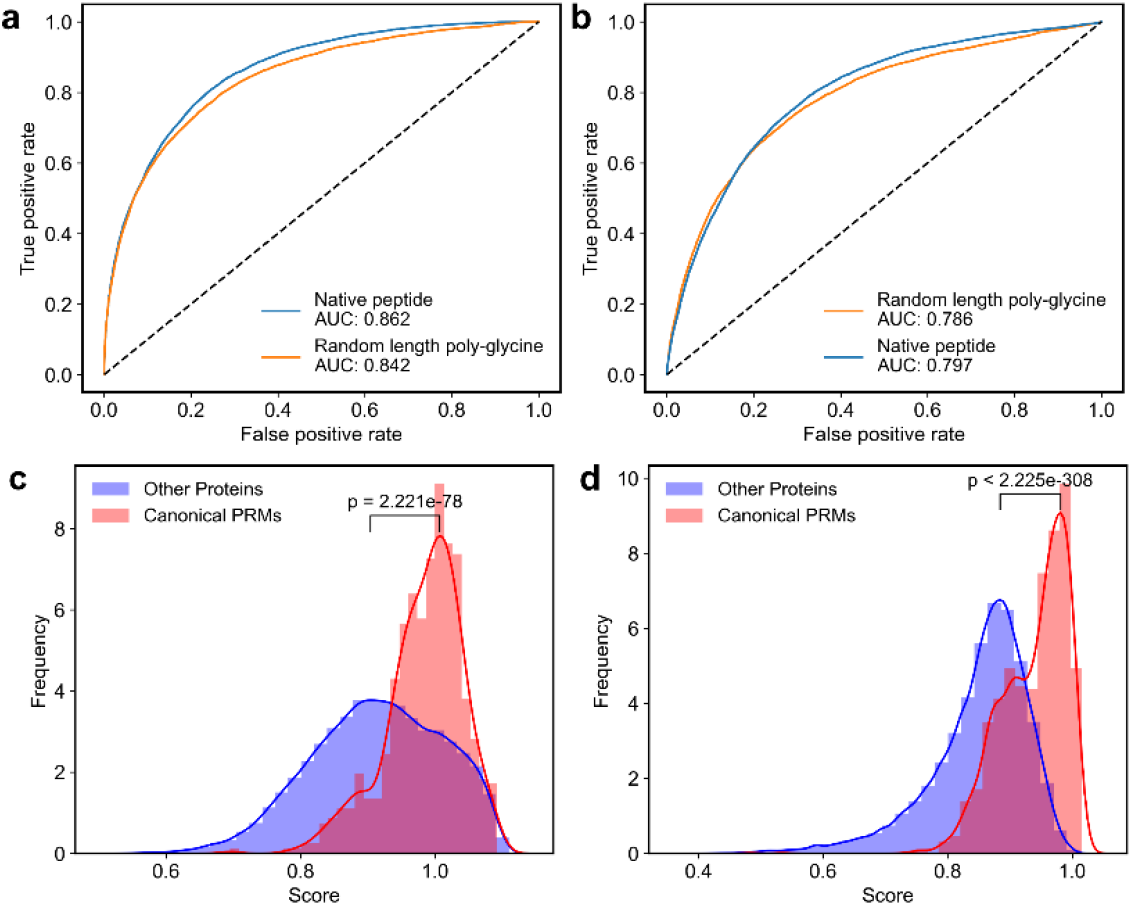
Peptide-agnostic binding site prediction using PepNN-Struct and PepNN-Seq. **a)** ROC curves on the validation dataset using PepNN-Struct with different input peptide sequences. **b)** ROC curves on the validation dataset using PepNN-Seq with different input peptide sequences. **c)** Scores assigned by PepNN-Struct to different domains in the PDB (Wilcoxon rank-sum test). **d)** Scores assigned by the PepNN-Seq to different domains in the reference human proteome (Wilcoxon rank-sum test).

To quantify the model’s confidence that a protein is a peptide-binding module, we generated a score that takes into account the binding probabilities that the model assigns the residues in the protein, as well as the percentage of residues that the model predicts are binding residues with high confidence. To compute this score, a Gaussian distribution was fit to the distribution of binding residue percentages in each protein from the training dataset (Fig S4a). The resulting score was the weighted average of the top *n* residue probabilities and the likelihood that a binding site would be composed of those *n* residues based on the aforementioned distribution. For each protein, *n* was chosen to maximize the score. As done in a previous study^11^, the weight assigned to each component of the score was chosen to maximize the correlation between the MCC of the prediction for each protein in the validation dataset, and its score (Fig S4b, c). This was motivated by the fact that the confidence of the model should correlate with its correctness.

We used the models to predict binding sites for domains in every unique chain in the PDB not within 30% homology of a sequence in the training dataset and domains in every sequence in the reference human proteome from UniProt^26^, not within 30% homology of a sequence in the training dataset. Domains were extracted by assigning PFAM^27^ annotations using InterProScan^28^ (Table S1, S2). To assess the capacity of the models to discriminate between peptide binding modules and other domains, we compared the distribution of scores for canonical PRMs to that of other proteins. Previously defined modular protein domains^29^, and peptide binding domains^3^ were considered canonical PRMs. In both the case of the PDB and the human proteome, the distribution of scores for canonical PRMs was higher than the background distribution (Fig 3c, d).

In total, PepNN-Struct assigns 39 623 domains in the PDB a score higher than the mean PRM score and PepNN-Seq assigns 10 332 domains in the human proteome a score higher than the mean PRM score. Analysis of the distribution of scores for different domains reveals that many DNA binding domains, including different transcription factors and DNA modifying enzymes, were assigned low scores on average by PepNN (Table S3, S4). This indicates that PepNN has the capacity to discriminate between different types of binding sites. There are, nonetheless, some nucleic acid binding domains with high scores (Table S3, S4) suggesting that there are false positives and that downstream computational and experimental work is required to validate putative peptide binding sties.

One domain identified by PepNN-Struct is the sterile alpha motif (SAM) domain of the Deleted-in-liver cancer 1 (DLC1) protein (Table S1). This domain was recently shown to be a peptide binding module^30^, demonstrating the capacity of the model to identify novel peptide binders. Another interesting hit identified using PepNN-Struct is the ORF7a accessory protein from the SARS-Cov-2 virus (Table S1). The model predicts that this protein has a peptide binding site located between two beta-sheets at the N-terminal end of the protein (Fig 4a). Validating this peptide binding site involves identifying a binding peptide and showing that the residues that comprise the binding site are necessary for the interaction. The ORF7a homolog from SARS-Cov has been shown to bind the ectodomain of the human BST-2 protein^31^. BST-2 binds and tethers viral particles to the cell membrane, thereby preventing viral exit^31^. It was shown that by binding BST-2, ORF7a prevents its glycosylation and thus reduces its ability to inhibit viral exit^31^. Given the fact that BST-2 forms a coiled-coil structure, it is possible that a linear segment along one of its helices binds to ORF7a at the predicted peptide-binding pocket.

**Figure 4:**
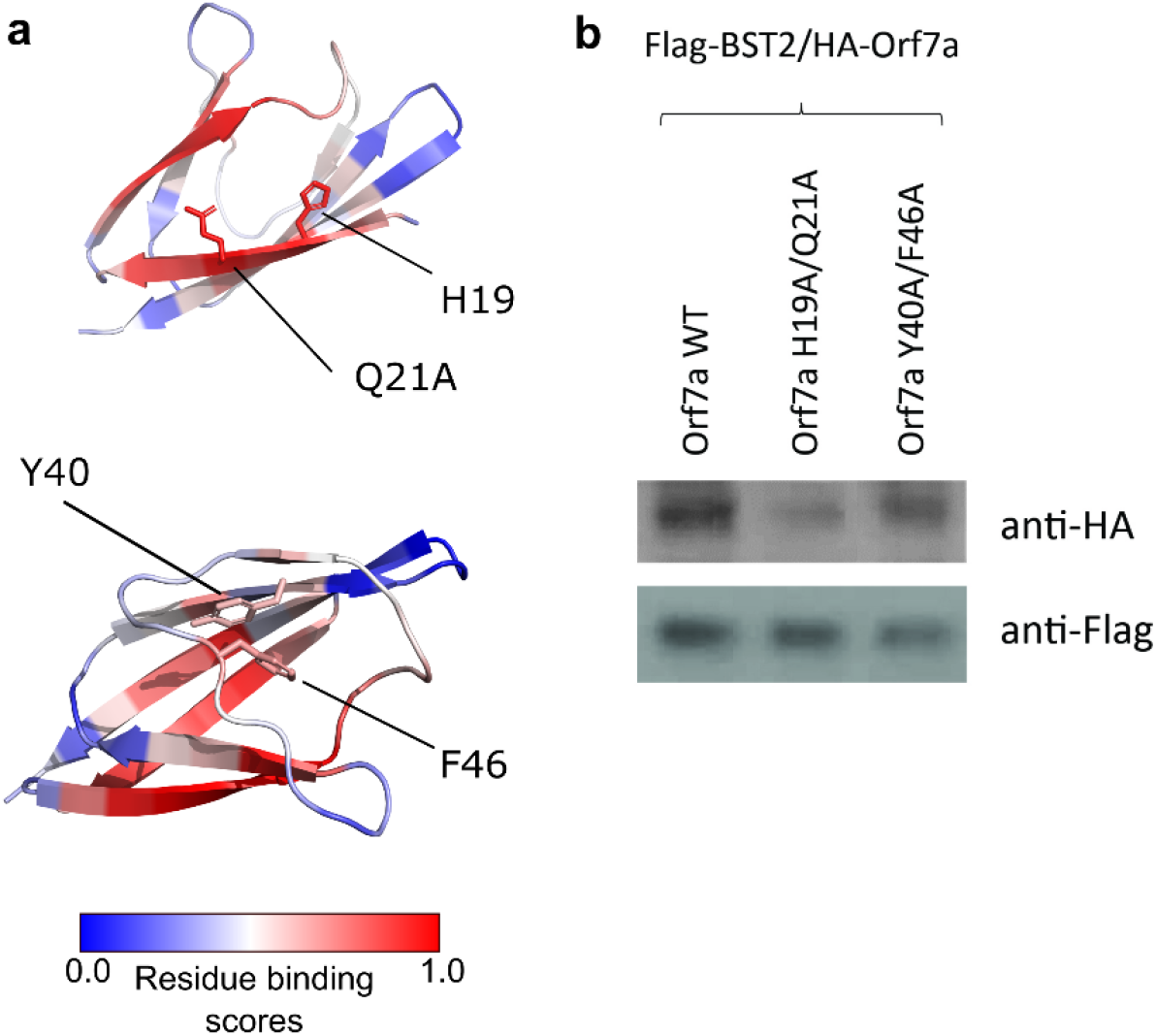
**a)** ORF7a peptide binding site prediction and key residues at the predicted binding site and an alternate binding site. **B)** Co-immunoprecipitation of wild type and mutant ORF7A with BST-2.

As a unbiased test of this prediction, we performed global docking of BST-2 onto ORF7a using the ClusPro webserver^32,33^. In seven of the top ten poses, BST-2 was found to interact with ORF7a at the predicted binding site (Fig S5). Based on the docking poses, alanine substitutions were introduced at two residues in the predicted binding site and two residues at an alternative binding site (Fig 4a). A co-immunoprecipitation assay demonstrated that the residues at the predicted binding site (H19A/Q21A) are necessary for ORF7a/BST-2 binding, corroborating the PepNN prediction (Fig 4b).

### Application of PepNN to epitope prediction

The binding of antibodies to their target antigens is largely facilitated by a set of variable segments known as complementarity-determining regions (CDRs). It has been shown that synthetic peptides derived from the sequences of these CDRs can bind the target antigen of the antibody from which they were derived^34–36^. We thus re-trained PepNN to predict the binding sites of different CDRs given an antigen structure. The estimated interface energy of peptide-protein complexes is greater than that of CDR-protein complexes (Fig S6). The pre-training dataset was consequently remade with less stringent thresholds (see Methods). We also ensured that fragments forming helix or strand secondary structures were filtered from the pre-training dataset. We trained the model to predict binding sites given H1, H2, and H3 loops. To generate a full epitope prediction, we assigned each residue the maximum score from the three models.

Overall, the observed performance was worse than that on peptide binding site prediction (Fig S7a). Nevertheless, the model makes reasonable predictions on numerous test antigens (Fig S7b).

## Discussion

We have developed parallel structure and sequence-based models for the prediction of peptide binding sites. These models, PepNN-Struct and PepNN-Seq, make use of a novel attention-based deep learning module that is integrated with transfer learning to compensate for the scarcity of peptide-protein complex data. Comparison to existing approaches shows that PepNN achieves state-of-the-art on the task of identifying peptide binding sites. In addition, unlike previously developed approaches, PepNN does not rely on structural or sequence alignments and is thus not dependent on the presence of structural data for homologs. Given the success of this approach, PepNN can be incorporated into local docking pipelines in order to facilitate the generation of protein-peptide complex models, a necessary step in delineating the molecular mechanisms underlying many cellular processes.

We furthermore demonstrated that PepNN can make accurate peptide-agnostic predictions. This observation is concordant with recent work that has suggested that a protein’s surface contains the majority of information regarding its capacity for biomolecular interactions ^37^. Other approaches, trained on negative binding data, are better suited than PepNN to discriminate between identified binding peptides^3,38^. By contrast, PepNN can uniquely be used to score proteins lacking a known peptide ligand to predict their ability to bind peptides. Running this procedure on all proteins in the PDB and the reference human proteome revealed a number of putative novel peptide recognition modules, suggesting that a large portion of the space of PRMs has yet to be characterized. As a demonstration of the model’s capacity to identify novel peptide binders, we showed that residues at a predicted peptide binding site are critical for the interaction between ORF7a and BST-2. The observation that PepNN can make predictions in the absence of a known peptide binder can also be used to discern regions of proteins that can be readily targeted by peptides. PepNN predictions can thus be used to inform the application of high-throughput experimental approaches to different proteins for the purpose of identifying therapeutic peptides. More generally, the success of PepNN serves as a proof-of-concept of the efficacy of reciprocal attention. This module can effectively be used to model bidirectional relationships between pairs of data points, and can thus be extended to other biomolecular interactions, including protein-protein and protein-DNA interactions. The use of transfer learning in the development of PepNN is also instructive for the development of approaches to solve related problems. The generated pre-training datasets of protein fragment-protein complexes are a valuable resource for modelling both peptide-protein and antibody-protein interactions.

## Materials and Methods

### Datasets

A dataset of protein-peptide complexes was generated by filtering complexes in the PDB. Crystal structures with a resolution of at least 2.5 □ that contain a chain of at least 50 amino acids in complex with a chain of 25 or less amino acids were considered putative peptide-protein complexes. Using FreeSASA^39^, complexes with a buried surface area of less than 400 □^2^ were filtered out, leaving 3046 complexes. The sequences of the receptors in the remaining complexes were clustered at a 30% identity threshold using PSI-CD-HIT^40^, and the resulting clusters were divided into training, validation, and test sets at proportions of 80%, 10% and 10% respectively. The test set contains 305 examples and is referred to as TS305.

A similar process was used to generate a dataset of protein fragment-protein complexes. Using the PeptiDerive Rosetta protocol^25^, the PDB was scanned for protein fragments of length 5-25 amino acids with a high predicted interface energy when in complex with another chain of at least 50 amino acids. Complexes were filtered out based on the distribution of predicted interface energies from the dataset of real protein-peptide complexes. Only complexes with an interface score less than one standard deviation above the mean of the peptide-protein complex distribution were maintained. The complexes were also filtered by buried surface area. Complexes with less than 400 □ ^2^ were once again filtered out. The final dataset contained 406 365 complexes. For data splitting, complexes were again clustered at 30% identity. In both datasets, binding residues were defined as those residues in the protein receptor with a heavy atom within 6 □ ^2^ from a heavy atom in the interacting chain.

In addition to TS305, the models were also tested on benchmark datasets compiled in other studies. This includes the test dataset used to evaluate the Interpep approach ^11^ (TS251), the test dataset used to evaluate the PepBind approach^12^ (TS639), and the test dataset used to evaluate SPRINT-Str ^41^ (TS125).

Complexes from the non-redundant SAbDab dataset were used for training the model to predict epitopes^42^. CDRs were defined using the PyIgClassify dataset^43^. Complexes where a particular CDR was not in contact with the antigen were filtered out when training the model to predict the binding site of that CDR. Antigen sequences were clustered at 30% identity before splitting the dataset. For pre-training on fragment-protein complexes, less stringent thresholds of a Rosetta interface score of -10 and a buried surface area of 250 □^2^ were used for filtering. In addition, secondary structure annotations were assigned to each fragment in the dataset using the MDTraj software^44^, and any fragment with more than two residues in the helix or strand classes were filtered out. The resulting dataset contained 684 912 entries.

### Input representation

In the case of PepNN-Struct, input protein structures are encoded using a previously described graph representation^19^, with the exception that additional node features are added to encode the side chain conformation at each residue. In this representation, a local coordinate system is defined at each residue based on the relative position of the Cα to the other backbone atoms^19^. The edges between residues encode information about the distance between the resides, the relative direction from one Cα to another, a quaternion representation of the rotation matrix between the local coordinate systems, and an embedding of the relative positions of the residues in the protein sequence^19^. The nodes include a one-hot representation of the amino acid identity and the torsional backbone angles^19^.

To encode information about the side-chain conformation, the centroid of the heavy side chain atoms at each residue is calculated. The direction of the atom centroid from the Cα is represented using a unit vector based on the defined local coordinate system. The distance is encoded using a radial basis function, similar to the encoding used for inter-residue distances in the aforementioned graph representation^19^. A one-hot encoding is used to represent protein and peptide sequence information. The pre-trained contextualized language model, ProtBert^23^, is used to embed the protein sequence in PepNN-Seq.

### Model architecture

The developed architecture takes inspiration the original Transformer architecture^18^, as well the Structured Transformer, developed for the design of proteins with a designated input structure^19^. Like these models, the PepNN architecture consists of repeating attention and feed forward layers (Fig 1a). PepNN differs from conventional Transformers, however, in that does not follow an encoder-decoder attention architecture and it makes use of multi-head reciprocal attention. This is a novel attention-based module that shares some conceptual similarity to a layer that was recently used for salient object detection^45^. Conventional scaled dot attention, mapping queries, represented by matrix *Q*, and key-value pairs, represented by matrices *K* and *V*, to attention values takes the following form^18^ :

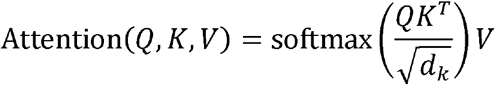

In reciprocal attention modules, protein residue embeddings are projected to a query matrix, 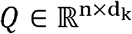 and a value matrix,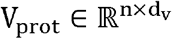 where *n* is the number of protein residues. Similarily, the peptide residue embeddings are projected a key matrix, 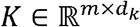 and a value matrix, 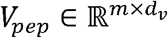, where *m* is the number of peptide residues. The resulting attention values are as follows:

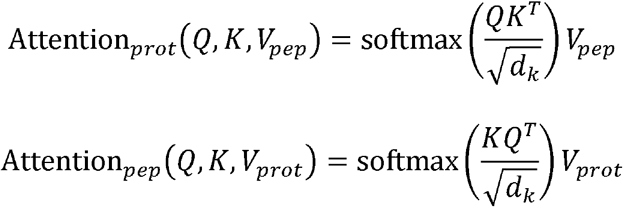

Projecting the residue encodings multiple times and concatenating the resulting attention values allows extension to multiple heads, as described previously^18^. The overall model architecture includes alternating self-attention and reciprocal attention layers, with a final set of layers to project the protein residue embedding down to a residue-wise probability score (Fig 1a). For the purpose of regularization, dropout layers were included after each attention layer.

Model hyperparameters were tuned using random search to optimize the cross-entropy loss on the fragment complex validation dataset. Specifically, eight hyperparameters were tuned; *d*_*model*_ (the model embedding dimension), *d*_*i*_ (the dimension of the hidden layer in the feed forward layers), *d*_*k*_, *d*_*v*_, the dropout percentage, the number of repititions of the reciprocal attention module, the number of heads in each attention layer, and the learning rate. In total, 100 random hyperparameter trials were attempted. *d*_*model*_ was set to 64, *d*_*i*_ was set to 64, *d*_*k*_ was set to 64, *d*_*v*_ was set to 128, dropout percentage was set to 0.2, the number of repetitions of the reciprocal attention module was set to 6, and each multi-head attention layer was composed of 6 heads.

### Training

Training was done using an Adam optimizer with a learning rate of 1e-4. A weighted cross-entropy loss was optimized to take into account the fact that the training dataset is skewed towards non-binding residues. In both the pre-training step with the fragment complex dataset and the training with the peptide complex dataset, early stopping was done based on the validation loss. Training was at most 500 000 iterations during the pre-training step and the at most 25 000 iterations during the fine-tuning step.

### Scoring potential novel peptide binding sites

Peptide-agnostic prediction of proteins in the human proteome and the PDB was performed by providing the model with a protein sequence/structure and a poly-glycine sequence of length 10 as the peptide. When computing scores using PepNN-Struct, the weight given to the top residue probabilities was 0.97. When computing scores using PepNN-Seq, this weight was set to 0.99. Pairwise comparisons were done with the distributions of every PFAM domain to remaining domains with a Wilcoxon rank-sum test and multiple testing correction was done using the Benjamini-Hochberg procedure.

### Statistical tests

Wilcoxon signed-rank and rank-sum tests were done using the SciPy python library^46^. Multiple testing correction was done using the statsmodels python package^47^. The DeLong test was done using the pROC R package^48^.

### Protein-protein docking of ORF7a/BST-2

The structure of the SARS-CoV-2 ORF7a encoded accessory protein (PDB ID 6W37) and mouse BST-2/Tetherin Ectodomain (PDB ID 3NI0^49^) were used as input structures for the ClusPro webserver^32,33^. The top ten poses, ranked by binding affinity, were used for downstream analysis.

### Cell lines and reagents

HEK293T cells were maintained in DMEM (ATCC) supplemented with 10% FBS and 1% pen/strep/glutamine, and the appropriate selection antibiotics when required. HA antibodies were obtained from Santa Cruz (7392) and Flag antibodies were purchased from Sigma (A8592).

### Western blotting

Transfected cells were scraped from 6-well dishes and lysed with lysis buffer (50 mM Tris-HCl pH7.4, 1% Nonidet P-40, 150 mM NaCl, 1 mM EDTA, 1× protease inhibitor mixture (Sigma)) for 30 min at 4 oC. The insoluble pellet was removed following a 10,000 rpm spin for 5 minutes at 4oC. Lysates were analyzed by SDS-PAGE/western blot using 4-20% Mini-PROTEAN Tris-glycine gels (Bio-Rad) transferred to PVDF membranes and blocked in 5% milk containing PBS-Tween-20 (0.1%) for 1 hour. PVDF membranes were then incubated with specified primary antibodies followed by incubation with horseradish peroxidase-conjugated secondary antibodies (Santa Cruz Biotechnology) and detected using enhanced chemiluminescence (GE Healthcare).

### Flag co-immunoprecipitation

HEK293T cells were cotransfected with Flag-tagged protein and HA-tagged protein. Cells were lysed 48 h after transfections with radioimmune precipitation assay buffer (50 mM Tris-HCl pH7.4, 1% Nonidet P-40, 150 mM NaCl, 1 mM EDTA, 10 mM Na3VO4, 10 mM sodium pyrophosphate, 25 mM NaF, 1× protease inhibitor mixture (Sigma)) for 30 min at 4 oC and coimmunoprecipitated with Flag beads (Clontech). The resulting immunocomplexes were analyzed by Western blot using the appropriate antibodies. Protein samples were separated using 4-20% Mini-PROTEAN Tris-glycine gels (Bio-Rad) transferred to PVDF membranes and blocked in 5% milk containing PBS-Tween-20 (0.1%) for 1 hour. PVDF membranes were then incubated with specified primary antibodies followed by incubation with horseradish peroxidase-conjugated secondary antibodies (Santa Cruz Biotechnology) and detected using enhanced chemiluminescence (GE Healthcare).

## Code and data availability

The datasets used in this study and the code to run PepNN are available at https://gitlab.com/oabdin/pepnn.

---

**Supplementary Figure 1:**
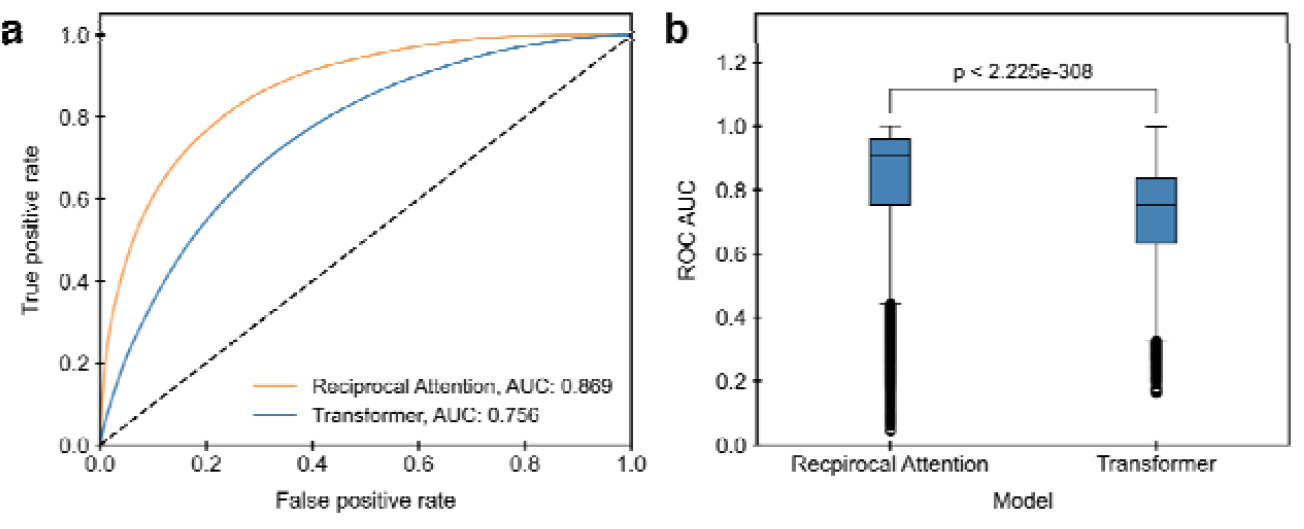
Comparison of the performance of the developed model to a Transformer with the same hyperparameters on the fragment complex validation dataset. **a)** ROC curves on all residues in the dataset. **b)** Comparison of distribution of ROC AUCs on different input proteins (Wilcoxon signed-rank test).

**Supplementary Figure 2:**
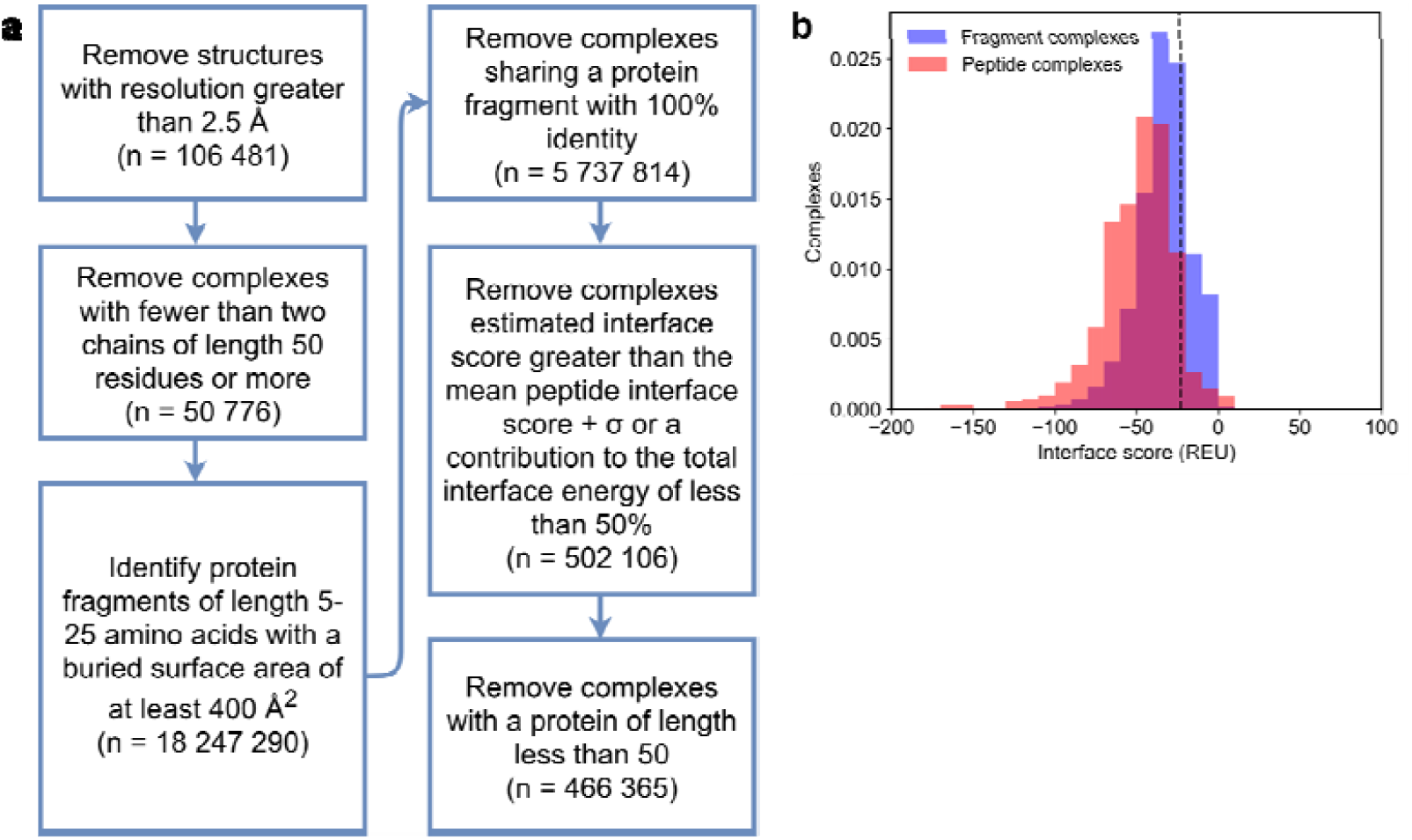
**a)** Curation pipeline for generation of a protein fragment-protein dataset. **b)** Comparison of estimated interface distribution for the all fragment-protein complexes and the dataset of peptide-protein complexes. The dashed lines indicate the threshold used for filtering fragment-protein complexes.

**Supplementary Figure 3:**
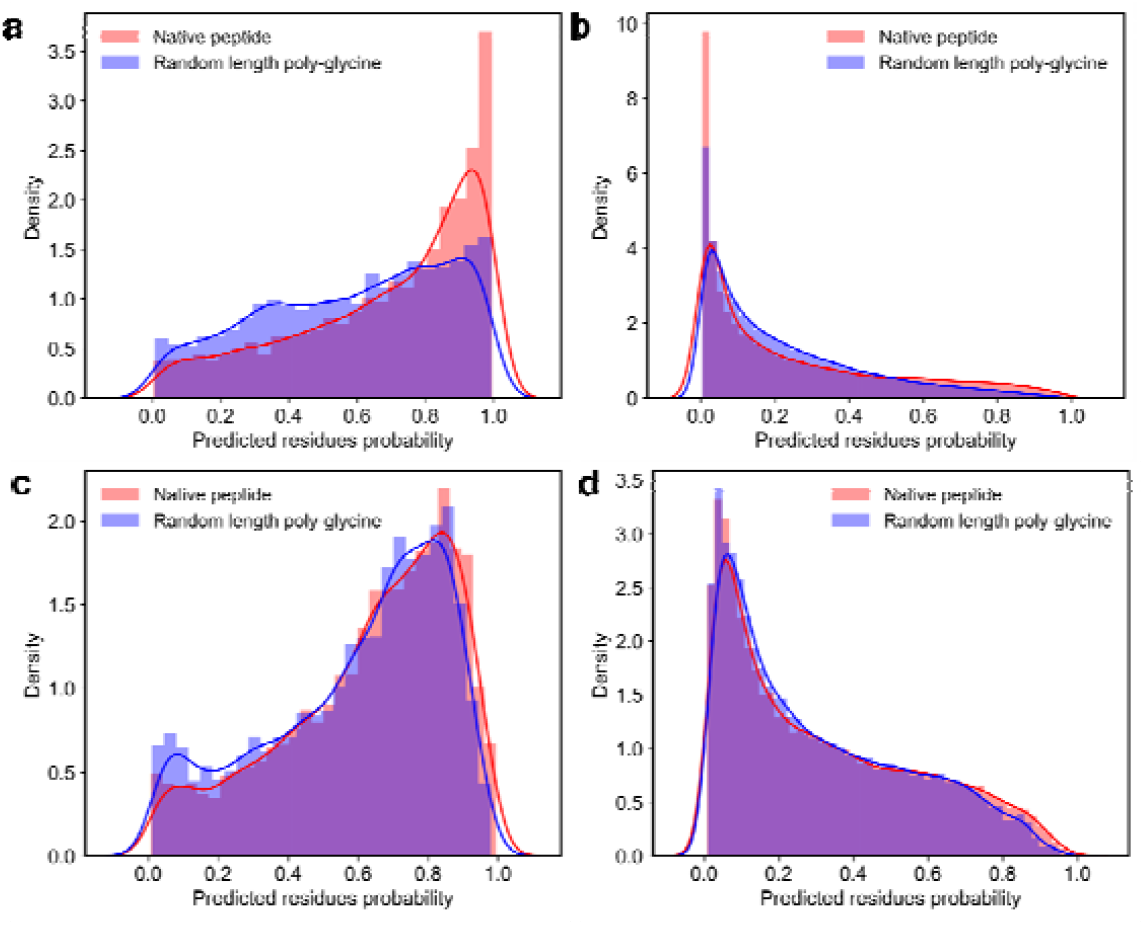
Probabilities assigned by PepNN-Struct and PepNN-Seq to different models residues with and without the native peptide sequence. **a)** Probabilities assigned by PepNN-Struct to binding residues. **b)** Probabilities assigned by PepNN-Struct to non-binding residues. **c)** Probabilities assigned by PepNN-Seq to binding residues. **d)** Probabilities assigned by PepNN-Seq to non-binding residues.

**Supplementary Figure 4:**
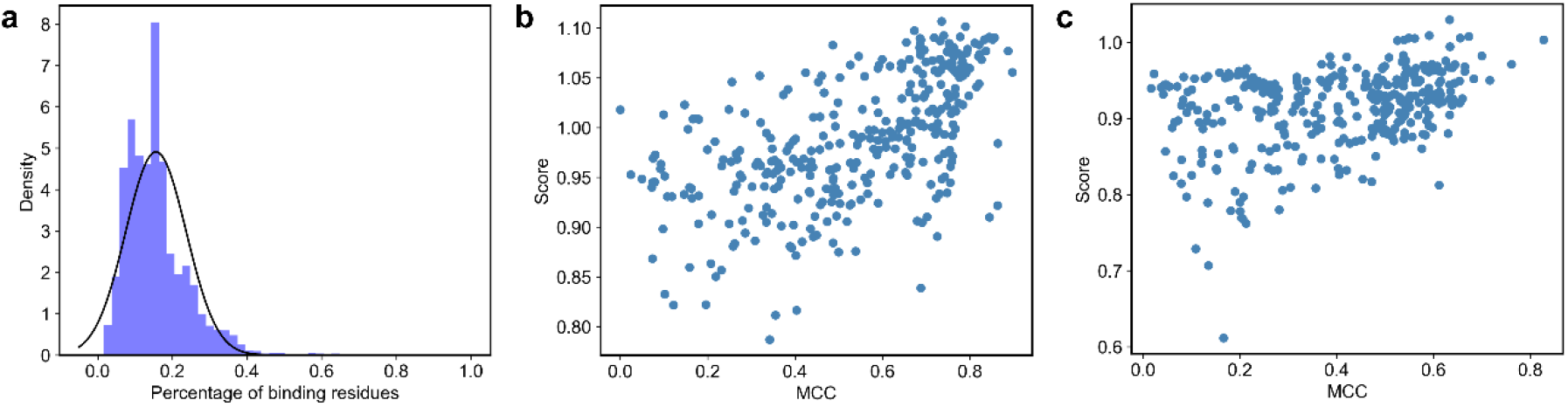
**a)** The percentage of binding residues in different examples in the training dataset. **b)** Relationship between scores assigned by PepNN-Struct and MCC of predictions. **c)** Relationship between scores assigned by PepNN-Seq and MCC of predictions.

**Supplementary Figure 5:**
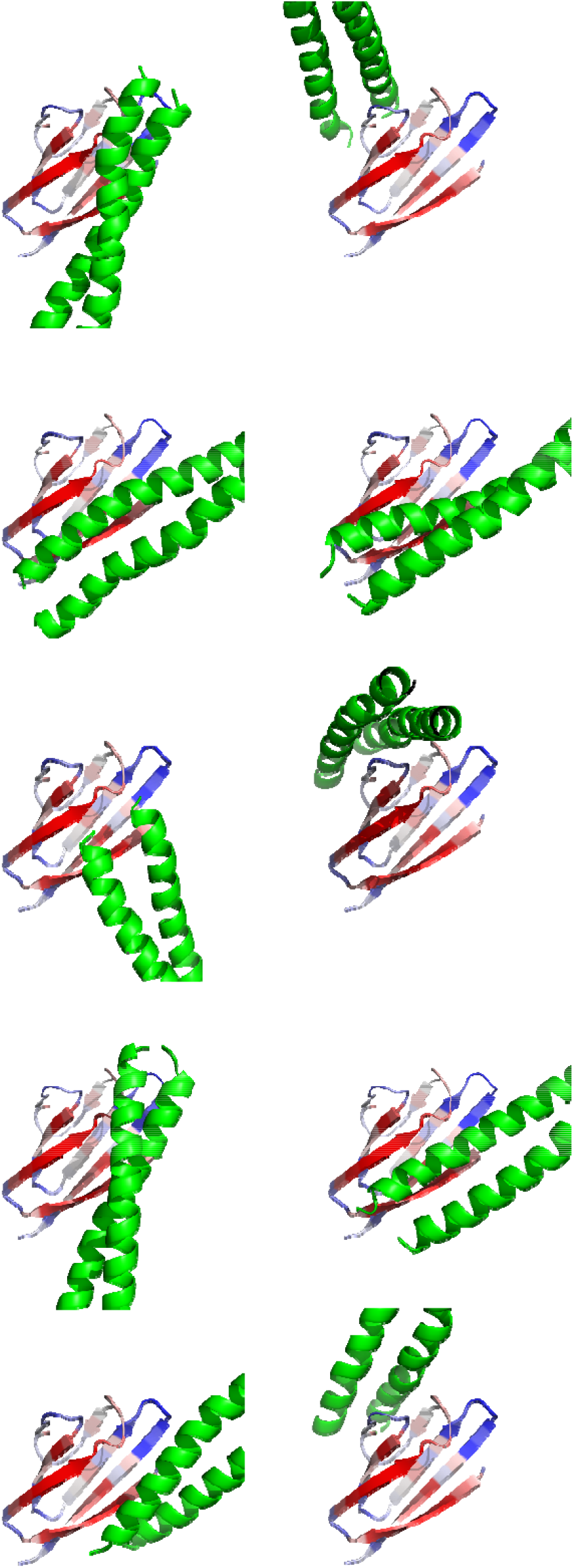
Complex conformations generated by docking BST-2 against ORF7a.

**Supplementary Figure 6:**
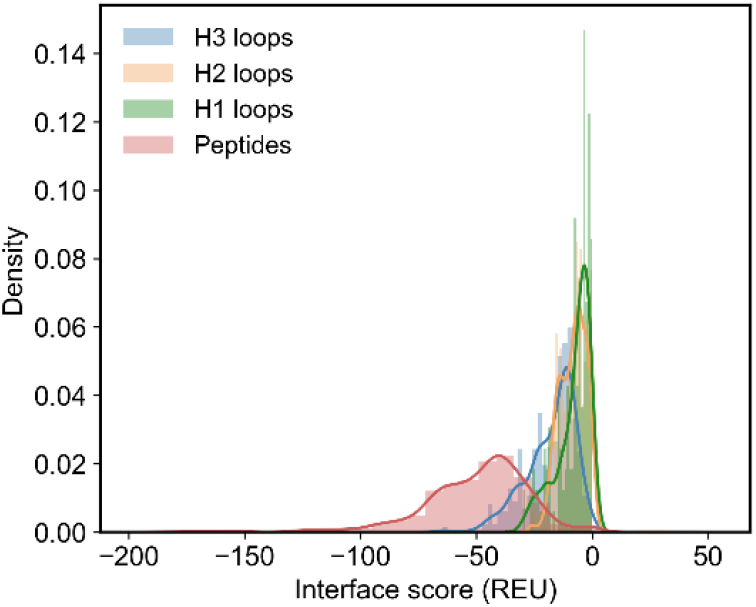
Comparison of estimated interface energies for peptide-protein complexes and CDR-protein complexes.

**Supplementary Figure 7:**
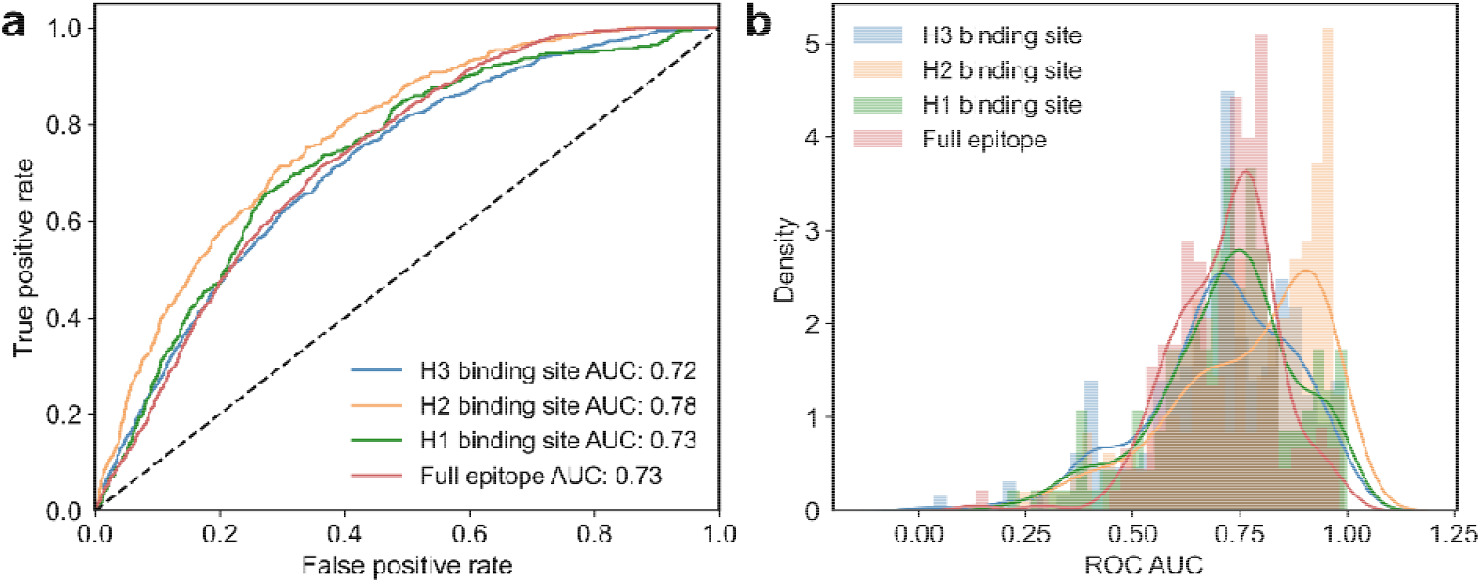
Performance of PepNN on the task of epitope prediction. **a)** ROC curves on all residues in the test dataset. **b)** Distribution of ROC AUC for different input antigens from the test set.

## References

1. Tompa, P., Davey, N. E., Gibson, T. J. & Babu, M. M. A Million peptide motifs for the molecular biologist. Molecular Cell (2014) doi:10.1016/j.molcel.2014.05.032.

2. Krumm, B. E. & Grisshammer, R. Peptide ligand recognition by G protein-coupled receptors. Front. Pharmacol. 6, 48 (2015).

3. Cunningham, J. M., Koytiger, G., Sorger, P. K. & AlQuraishi, M. Biophysical prediction of protein–peptide interactions and signaling networks using machine learning. Nat. Methods 17, 175–183 (2020).

4. Yang, F. et al. Protein Domain-Level Landscape of Cancer-Type-Specific Somatic Mutations. PLOS Comput. Biol. 11, 1–30 (2015).

5. Hagai, T., Azia, A., Babu, M. M. & Andino, R. Use of Host-like Peptide Motifs in Viral Proteins Is a Prevalent Strategy in Host-Virus Interactions. Cell Rep. 7, 1729–1739 (2014).

6. Ciemny, M. et al. Protein–peptide docking: opportunities and challenges. Drug Discovery Today (2018) doi:10.1016/j.drudis.2018.05.006.

7. Raveh, B., London, N. & Schueler-Furman, O. Sub-angstrom modeling of complexes between flexible peptides and globular proteins. Proteins Struct. Funct. Bioinforma. 78, 2029–2040 (2010).

8. London, N., Raveh, B. & Schueler-Furman, O. Modeling Peptide--Protein Interactions. in Homology Modeling: Methods and Protocols (eds. Orry, A. J. W. & Abagyan, R.) 375–398 (Humana Press, 2012). doi:10.1007/978-1-61779-588-6_17.

9. Agrawal, P. et al. Benchmarking of different molecular docking methods for protein-peptide docking. BMC Bioinformatics 19, 426 (2019).

10. Weng, G. et al. Comprehensive Evaluation of Fourteen Docking Programs on Protein– Peptide Complexes. J. Chem. Theory Comput. 16, 3959–3969 (2020).

11. Johansson-Åkhe, I., Mirabello, C. & Wallner, B. Predicting protein-peptide interaction sites using distant protein complexes as structural templates. Sci. Rep. 9, (2019).

12. Zhao, Z., Peng, Z. & Yang, J. Improving Sequence-Based Prediction of Protein-Peptide Binding Residues by Introducing Intrinsic Disorder and a Consensus Method. J. Chem. Inf. Model. 58, (2018).

13. Taherzadeh, G., Yang, Y., Zhang, T., Liew, A. W.-C. & Zhou, Y. Sequence-based prediction of protein–peptide binding sites using support vector machine. J. Comput. Chem. 37, 1223–1229 (2016).

14. Taherzadeh, G., Zhou, Y., Liew, A. W. C. & Yang, Y. Structure-based prediction of protein-peptide binding regions using Random Forest. Bioinformatics 34, (2018).

15. Wardah, W. et al. Predicting protein-peptide binding sites with a deep convolutional neural network. J. Theor. Biol. 496, (2020).

16. Iqbal, S. & Hoque, M. T. PBRpredict-Suite: a suite of models to predict peptide-recognition domain residues from protein sequence. Bioinformatics 34, 3289–3299 (2018).

17. Senior, A. W. et al. Improved protein structure prediction using potentials from deep learning. Nature 577, 706–710 (2020).

18. Vaswani, A. et al. Attention is all you need. in Advances in Neural Information Processing Systems (2017).

19. Ingraham, J., Garg, V. K., Barzilay, R. & Jaakkola, T. Generative models for graph-based protein design. in Deep Generative Models for Highly Structured Data, DGS@ICLR 2019 Workshop (2019).

20. Strokach, A., Becerra, D., Corbi-Verge, C., Perez-Riba, A. & Kim, P. M. Fast and Flexible Protein Design Using Deep Graph Neural Networks. Cell Syst. 11, 402-411.e4 (2020).

21. Mohan, A. et al. Analysis of Molecular Recognition Features (MoRFs). J. Mol. Biol. 362, (2006).

22. London, N., Raveh, B., Movshovitz-Attias, D. & Schueler-Furman, O. Can self-inhibitory peptides be derived from the interfaces of globular protein-protein interactions? Proteins Struct. Funct. Bioinforma. (2010) doi:10.1002/prot.22785.

23. Elnaggar, A. et al. ProtTrans: Towards Cracking the Language of Life{\textquoteright}s Code Through Self-Supervised Deep Learning and High Performance Computing. bioRxiv (2020) doi:10.1101/2020.07.12.199554.

24. Rao, R. et al. Evaluating Protein Transfer Learning with {TAPE}. CoRR abs/1906.0, (2019).

25. Sedan, Y., Marcu, O., Lyskov, S. & Schueler-Furman, O. Peptiderive server: derive peptide inhibitors from protein-protein interactions. Nucleic Acids Res. (2016) doi:10.1093/nar/gkw385.

26. Consortium, T. U. UniProt: a worldwide hub of protein knowledge. Nucleic Acids Res. 47, D506–D515 (2018).

27. Finn, R. D. et al. Pfam: the protein families database. Nucleic Acids Res. 42, D222–D230 (2013).

28. Jones, P. et al. InterProScan 5: Genome-scale protein function classification. Bioinformatics (2014) doi:10.1093/bioinformatics/btu031.

29. Jadwin, J. A., Ogiue-Ikeda, M. & Machida, K. The application of modular protein domains in proteomics. FEBS Lett. 586, 2586–2596 (2012).

30. Joshi, R. et al. {D}{L}{C}1 {S}{A}{M} domain-binding peptides inhibit cancer cell growth and migration by inactivating {R}ho{A}. J Biol Chem 295, 645–656 (2020).

31. Taylor, J. K. et al. Severe Acute Respiratory Syndrome Coronavirus ORF7a Inhibits Bone Marrow Stromal Antigen 2 Virion Tethering through a Novel Mechanism of Glycosylation Interference. J. Virol. 89, 11820–11833 (2015).

32. Kozakov, D. et al. The ClusPro web server for protein--protein docking. Nat. Protoc. 12, 255–278 (2017).

33. Vajda, S. et al. New additions to the ClusPro server motivated by CAPRI. Proteins 85, 435–444 (2017).

34. Williams, W. V., Kieber-Emmons, T., VonFeldt, J., Greene, M. I. & Weiner, D. B. Design of bioactive peptides based on antibody hypervariable region structures: Development of conformationally constrained and dimeric peptides with enhanced affinity. J. Biol. Chem. 266, (1991).

35. Williams, W. V et al. Sequences of the cell-attachment sites of reovirus type 3 and its anti-idiotypic/antireceptor antibody: modeling of their three-dimensional structures. Proc. Natl. Acad. Sci. 85, 6488–6492 (1988).

36. Taub, R. et al. A monoclonal antibody against the platelet fibrinogen receptor contains a sequence that mimics a receptor recognition domain in fibrinogen. J. Biol. Chem. 264, (1989).

37. Gainza, P. et al. Deciphering interaction fingerprints from protein molecular surfaces using geometric deep learning. Nat. Methods 17, (2020).

38. Lei, Y. et al. CAMP: a Convolutional Attention-based Neural Network for Multifaceted Peptide-protein Interaction Prediction. bioRxiv (2020) doi:10.1101/2020.11.16.384784.

39. Mitternacht, S. FreeSASA: An open source C library for solvent accessible surface area calculations. F1000Research (2016) doi:10.12688/f1000research.7931.1.

40. Fu, L., Niu, B., Zhu, Z., Wu, S. & Li, W. CD-HIT: Accelerated for clustering the next-generation sequencing data. Bioinformatics (2012) doi:10.1093/bioinformatics/bts565.

41. Taherzadeh, G., Zhou, Y., Liew, A. W.-C. & Yang, Y. Structure-based prediction of protein– peptide binding regions using Random Forest. Bioinformatics 34, 477–484 (2017).

42. Dunbar, J. et al. SAbDab: the structural antibody database. Nucleic Acids Res. 42, D1140– D1146 (2014).

43. Adolf-Bryfogle, J., Xu, Q., North, B., Lehmann, A. & Dunbrack Jr, R. L. PyIgClassify: a database of antibody CDR structural classifications. Nucleic Acids Res. 43, D432–D438 (2015).

44. McGibbon, R. T. et al. MDTraj: A Modern Open Library for the Analysis of Molecular Dynamics Trajectories. Biophys. J. 109, 1528–1532 (2015).

45. Xia, C., Li, J., Su, J. & Tian, Y. Exploring Reciprocal Attention for Salient Object Detection by Cooperative Learning. (2019).

46. Virtanen, P. et al. SciPy 1.0: fundamental algorithms for scientific computing in Python. Nat. Methods 17, (2020).

47. Seabold, S. & Perktold, J. Statsmodels: econometric and statistical modeling with Python. in 9th Python in Science Conference (2010).

48. Robin, X. et al. pROC: an open-source package for R and S+ to analyze and compare ROC curves. BMC Bioinformatics 12, 77 (2011).

49. Swiecki, M. et al. Structural and biophysical analysis of BST-2/tetherin ectodomains reveals an evolutionary conserved design to inhibit virus release. J. Biol. Chem. 286, 2987–2997 (2011).

